## Supplementary Material for "Nutritional state-dependent modulation of Insulin-Producing Cells in *Drosophila*"

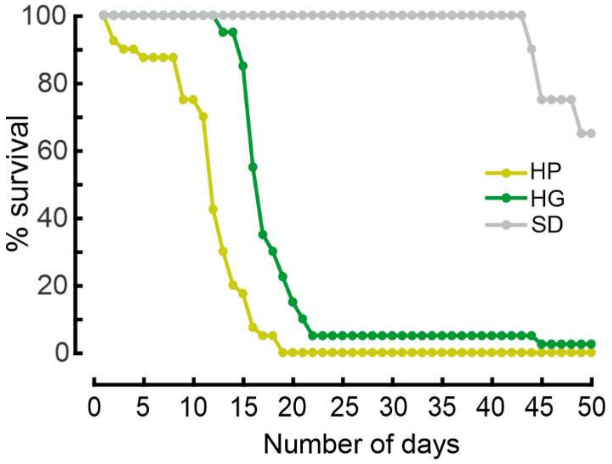


**Figure S1: Dietary restriction impairs survival in *Drosophila*.**

Percentage survival of flies on different diets. After 24 h of starvation, flies were kept on a high glucose (HG), high protein (HP) or standard diet (SD) and survival was scored every day. Data points were combined from two replicates of 20 flies per condition (N = 40 flies per condition total).

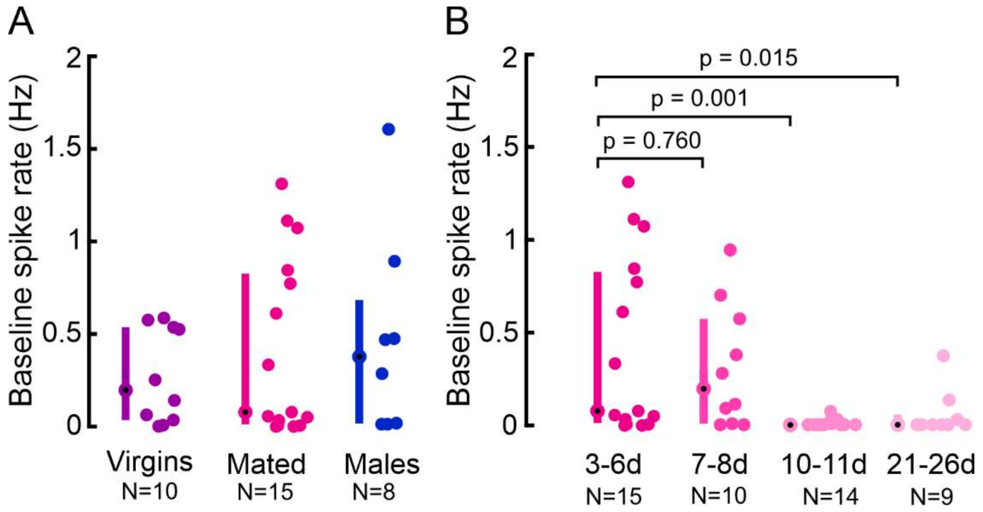


**Figure S2: Sex and mating state do not affect IPC activity, but aging does.**

**A)** Comparison of IPC baseline spike rate in virgin females, mated females, and males. **B)** Comparison of IPC baseline spike rate in different age groups. d = days, N = number of IPC recordings (see Table S5 for number of flies). Each dot represents one IPC, error bars indicate the median (circle) and interquartile range (bars). p-values calculated via Wilcoxon rank-sum test.

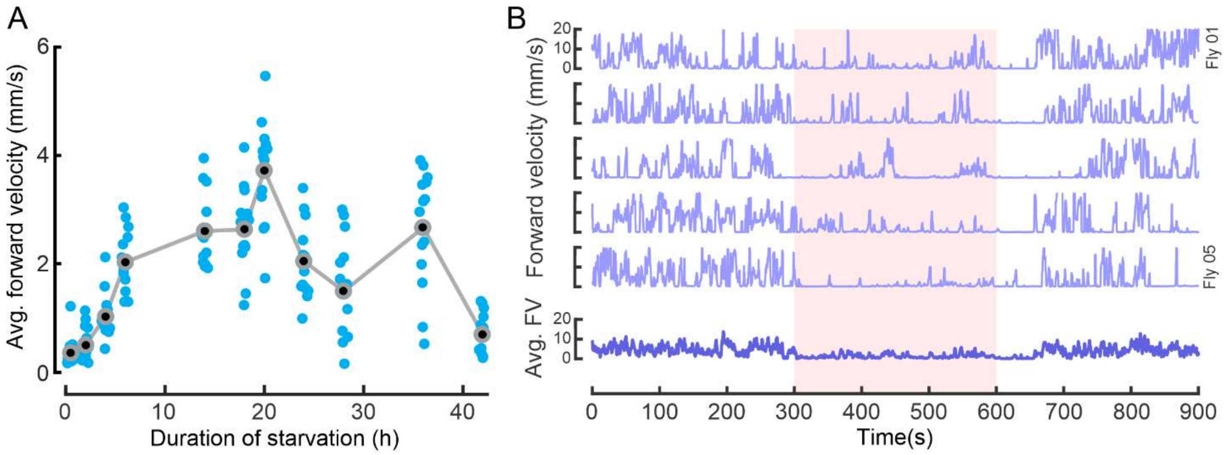


**Figure S3: Starvation duration and OAN activation affect locomotor activity.**

**A)** Average forward velocity of flies during different periods of starvation. N = 20 flies per condition, except for 14 h (N = 19), 20 h (N = 19) and 28 h (N = 16). Each point represents one fly, gray circles and lines represent means. **B)** Upper panel: Examples for forward velocities of five flies displaying stopping behavior during OAN activation (fifth activation cycle). Lower panel: Average forward velocity of all flies from one replicate (N = 20). Red bar represents OAN activation.

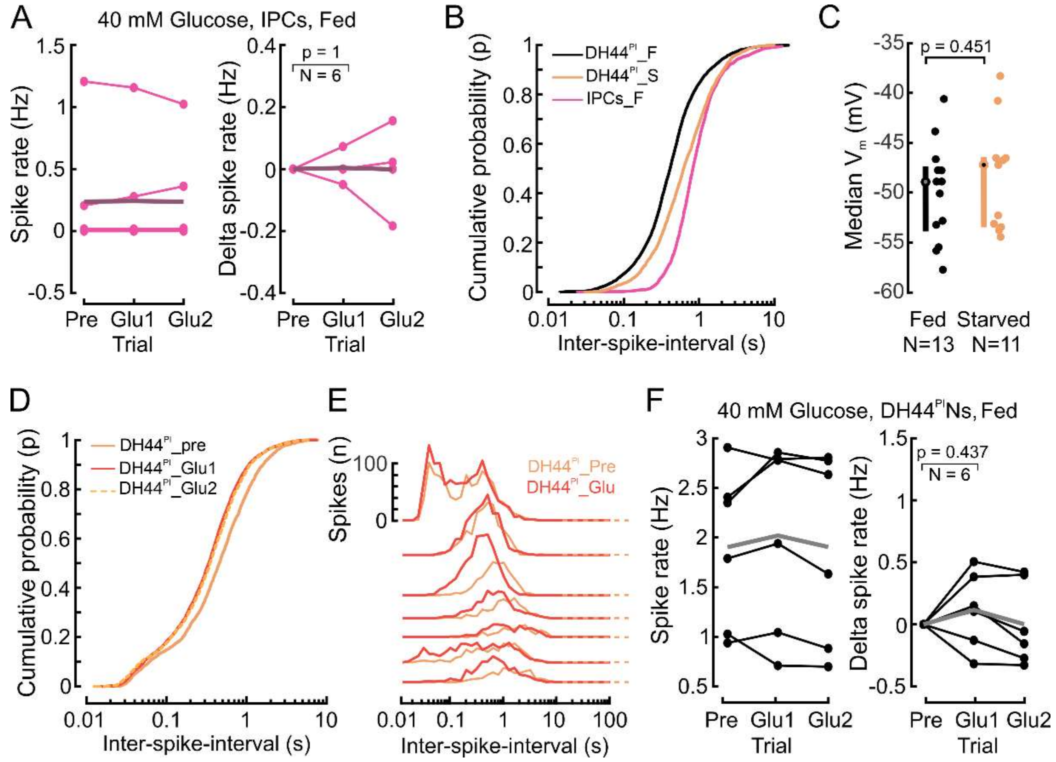


**Figure S4: Glucose perfusion does not affect the activity of IPCs and DH44$^{PI}$Ns in fed flies but shifts interspike interval distributions in DH44$^{PI}$Ns.**

**A)** IPC spike rate (left) and change in spike rate (right) in fed flies under 40 mM glucose perfusion. Pre, five-minute recording in glucose-free extracellular saline, Glu1 and Glu2, two subsequent five-minute recordings in glucose-rich extracellular saline, starting eight minutes after onset of glucose perfusion. Points represent mean of individual flies, thick line represents the grand mean. p-values were calculated via Wilcoxon signed-rank test. Delta spike rates were calculated by subtracting the mean spike rate during the Pre window. **B)** DH44$^{PI}$N spike activity patterns change with the nutritional state. Cumulative probability distribution of the ISI in DH44$^{PI}$Ns and IPCs. F = fed, S = starved flies. Distributions were compared using a two-sample Kolgomorov-Smirnov test (DH44$^{PI}$_F vs IPCs_F: p = 1.5e$^{-216}$; DH44$^{PI}$_F vs DH44$^{PI}$_S: p = 2.5e$^{-86}$; DH44$^{PI}$_S vs IPCs_F: p = 7.4e$^{-51}$). IPC dataset used in these panels corresponds to IPC baseline activity in fed flies in Figure 1D (N = 15). DH44$^{PI}$ dataset corresponds to DH44$^{PI}$N baseline activity in fed (N = 13) and starved flies (N = 11) in Figure 3F. **C)** Median membrane potential of DH44$^{PI}$Ns in fed and starved flies. **D)** Cumulative probability distribution of the DH44 ISI in Pre, Glu1 and Glu2 windows reveals that DH44$^{PI}$N activity became more bursty during glucose perfusion. Distributions were compared using a two-sample Kolgomorov-Smirnov test (DH44$^{PI}$_pre vs DH44$^{PI}$_Glu1: p = 6.5e$^{-30}$; DH44$^{PI}$_pre vs DH44$^{PI}$_Glu2: p = 2.3e$^{-21}$; DH44$^{PI}$_Glu1 vs DH44$^{PI}$_Glu2: p = 0.05, N = 7). **E)** DH44$^{PI}$N spike activity patterns change during glucose perfusion. ISIs are shown for DH44$^{PI}$Ns before (Pre) and during (Glu1 and Glu2) 40 mM glucose perfusion. **F)** DH44$^{PI}$N spike rate (left) and change in spike rate (right) in fed flies under 40 mM glucose perfusion. Plot details as in A.

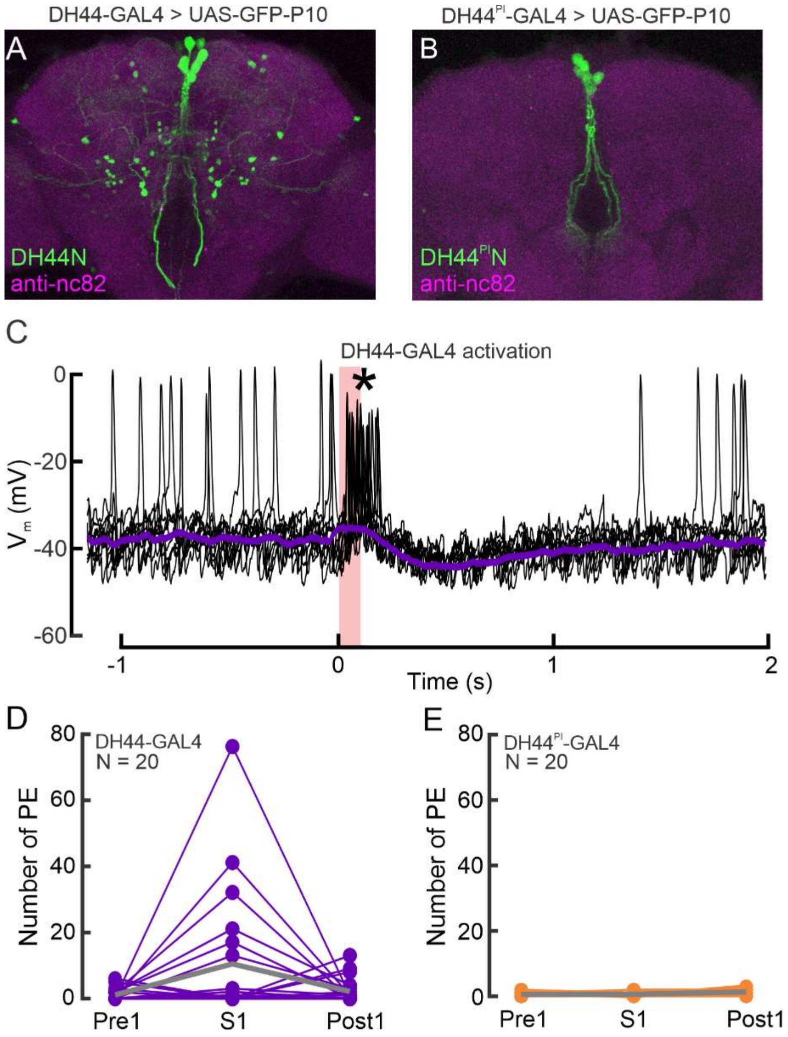


**Figure S5: Effects of differential activation of DH44Ns and DH44$^{PI}$Ns on IPCs and behavior.**

**A)** Expression pattern of the broad DH44N driver line in the brain visualized with a *UAS-GFP-P10* reporter. GFP was enhanced with anti-GFP (green), brain neuropils were stained with anti-nc82 (magenta). **B)** Expression pattern of the sparse DH44$^{PI}$N driver line. Staining details as in A. **C)** Example recording of an IPC during optogenetic activation of the broad DH44N driver line. In this one example, we observed strong activation of the IPC during DH44N activation (asterisk). About 100 ms after activation, the IPC was inhibited similar to all other recordings (see Figure 4B). **D)** Number of proboscis extensions (PE) before (Pre1), during (S1), and after the first LED pulse (Post1) activating DH44Ns in freely walking flies in the UFO. PEs were scored manually. Gray lines represent the grand mean. **E)** Activation of DH44$^{PI}$Ns did not drive PE. Plot details as in D.

**Table S1: p-values values for statistical comparisons in Figure 2, activation window.**

Comparisons were made between the respective driver lines and Empty-split control flies. p-values were determined using the Wilcoxon rank-sum test. Columns 1-5 represent the five optogenetic activation cycles in Figure 2E, H and K. 'Activation' shows the p-values comparing average forward velocity pooled across all activation trials (Figure 2L).

| Genotype | 1 | 2 | 3 | 4 | 5 | Activation |
|---|---|---|---|---|---|---|
| DILP2>CsChr, Fed | **0.0256** | 0.8694 | 0.5705 | 0.1867 | 0.1507 | **0.0133** |
| DILP2>CsChr, Starved | **$3.4e^{-09}$** | **$7.2e^{-06}$** | **$1.2e^{-06}$** | **0.0021** | **0.0042** | **$1.0e^{-20}$** |
| TDC2>CsChr, Fed | **$2.1e^{-05}$** | **$1.3e^{-10}$** | **$7.7e^{-12}$** | **$8.3e^{-12}$** | **$3.6e^{-12}$** | **$6.8e^{-48}$** |

**Table S2: p-values values for statistical comparisons in Figure 2, post activation window.**

P1-P5 represent the 'post activation' windows in Figure 2E, H and K. 'Post' contains the p-values comparing average forward velocity pooled across all trials (Figure 2M). Other details as for Table S1.

| Genotype | P1 | P2 | P3 | P4 | P5 | Post |
|---|---|---|---|---|---|---|
| DILP2>CsChr, Fed | 0.4357 | 0.9958 | 0.2543 | 0.7868 | 0.3874 | 0.6170 |
| DILP2>CsChr, Starved | **$5.3e^{-06}$** | **0.0003** | **0.0008** | 0.2322 | 0.4540 | **$1.7e^{-8}$** |
| TDC2>CsChr, Fed | **$6.5e^{-13}$** | **$1.7e^{-13}$** | **$8.9e^{-13}$** | **$4.7e^{-13}$** | **$7.2e^{-12}$** | **$7.3e^{-58}$** |

**Table S3: p-values values for statistical comparisons in Figure 4, activation window.**

1 - 5 represent the five activation cycles in Figure 4K and N. Other details as for Table S1.

| Genotype | 1 | 2 | 3 | 4 | 5 |
|---|---|---|---|---|---|
| DH44>CsChr | **$1.9e^{-11}$** | **$2.1e^{-12}$** | **$8.6e^{-08}$** | **$4.5e^{-06}$** | **$2.8e^{-06}$** |
| $DH44^{PI}$>CsChr | **0.0001** | **0.0022** | 0.0540 | **0.0156** | **0.0015** |

**Table S4: p-values values for statistical comparisons in Figure 4, post activation window.**

P1-P5 represent the 'post activation' windows in Figure 4K and N. Other details as for Table S1.

| Genotype | P1 | P2 | P3 | P4 | P5 |
|---|---|---|---|---|---|
| DH44>CsChr | **$2.3e^{-10}$** | **$2.7e^{-13}$** | **$2.8e^{-10}$** | **$4.0e^{-08}$** | **$6.7e^{-10}$** |
| $DH44^{PI}$>CsChr | 0.2711 | 0.1895 | 0.9044 | 0.0602 | **0.0054** |

**Table S5: Number of IPCs and number of flies in each experiment**.

*The total number of flies and IPCs is not equal to the sum of all datasets because 19 IPCs from 19 flies were used in two figures: The ‘Pre’ datasets from Figure 3A, 3B and S4A were also used to establish the population averages in Figure 1D, and the ‘Fed’ dataset from Figure 1D was reused for ‘Mated’ and ‘3-6d’ datasets in Figure S2.

| Dataset | Number of IPCs (N) | Number of flies |
|---|---|---|
| Figure 1D, Fed | 15 | 15 |
| Figure 1D, Starved | 23 | 19 |
| Figure 1G, 0.5-2h | 16 | 8 |
| Figure 1G, 6-8h | 10 | 7 |
| Figure 1G, HG, 3-5h | 7 | 4 |
| Figure 1G, HG, 6-12h | 11 | 9 |
| Figure 1G, HG, 18-24h | 11 | 9 |
| Figure 1H, HG+SD | 12 | 8 |
| Figure 1H, HF | 10 | 5 |
| Figure 1H, HA+SD | 10 | 5 |
| Figure 1H, SD | 12 | 7 |
| Figure 1H, HP | 11 | 9 |
| Figure 3A | 6 | 6 |
| Figure 3B | 7 | 7 |
| Figure 4C and 4D | 10 | 10 |
| Figure 4G and 4H | 8 | 5 |
| Figure S2A, Virgins | 10 | 9 |
| Figure S2A, Mated | 15 | 15 |
| Figure S2A, Male | 8 | 7 |
| Figure S2B, 3-6d | 15 | 15 |
| Figure S2B, 7-8d | 10 | 8 |
| Figure S2B, 10-11d | 14 | 8 |
| Figure S2B, 21-26d | 9 | 8 |
| Figure S4A | 6 | 6 |
| **Total** | **217*** | **160*** |

**Table S6: Number of $DH44^{PI}Ns$ in Figure 3.**

*The total number of flies and $DH44^{PI}Ns$ is not equal to the sum of individual datasets because the ‘Pre’ datasets from Figure 3G and Figure S4F were also used to establish population averages in Figure 3F.

| Dataset | Number of $DH44^{PI}Ns$ (N) | Number of flies |
|---|---|---|
| Figure 3F, Fed | 13 | 11 |
| Figure 3F, Starved | 11 | 10 |
| Figure 3G | 7 | 7 |
| Figure S4F | 6 | 6 |
| **Total** | **24*** | **21*** |

**Supplementary Video S1: Behavioral effects of optogenetic OAN activation on *Drosophila* locomotor activity in the UFO.**

Example video of OAN-activated flies walking in the UFO during the fifth activation cycle. The white box indicates when the optogenetic stimulation LED is on. The video includes one-minute before activation (P4 in Fig 2K), 5-minutes of activation (5 in Fig 2K), and one-minute after activation (P5 in Fig 2K). During OAN activation, the forward velocity decreased significantly, including several halting episodes.

**Supplementary Video S2: Behavioral effects of the optogenetic activation protocol on the locomotor activity of Empty split-Gal4 control flies in the UFO.**

Example video of control flies responding to the light pulse used for optogenetic activation during the fifth activation cycle. These flies were recorded in parallel to the OAN-activated flies in Video S1. Details as for Supplementary video S1.
